# YASS: Yet Another Spike Sorter

**DOI:** 10.1101/151928

**Authors:** JinHyung Lee, David Carlson, Hooshmand Shokri, Weichi Yao, Georges Goetz, Espen Hagen, Eleanor Batty, EJ Chichilnisky, Gaute Einevoll, Liam Paninski

## Abstract

Spike sorting is a critical first step in extracting neural signals from large-scale electrophysiological data. This manuscript describes an efficient, reliable pipeline for spike sorting on dense multi-electrode arrays (MEAs), where neural signals appear across many electrodes and spike sorting currently represents a major computational bottleneck. We present several new techniques that make dense MEA spike sorting more robust and scalable. Our pipeline is based on an efficient multi-stage “triage-then-cluster-then-pursuit” approach that initially extracts only clean, high-quality waveforms from the electrophysiological time series by temporarily skipping noisy or “collided” events (representing two neurons firing synchronously). This is accomplished by developing a neural network detection method followed by efficient outlier triaging. The clean waveforms are then used to infer the set of neural spike waveform templates through nonparametric Bayesian clustering. Our clustering approach adapts a “coreset” approach for data reduction and uses efficient inference methods in a Dirichlet process mixture model framework to dramatically improve the scalability and reliability of the entire pipeline. The “triaged” waveforms are then finally recovered with matching-pursuit deconvolution techniques. The proposed methods improve on the state-of-the-art in terms of accuracy and stability on both real and biophysically-realistic simulated MEA data. Furthermore, the proposed pipeline is efficient, learning templates and clustering much faster than real-time for a ≃ 500-electrode dataset, using primarily a single CPU core.

## 1 Introduction

The analysis of large-scale multineuronal spike train data is crucial for current and future neuroscience research. These analyses are predicated on the existence of reliable and reproducible methods that feasibly scale to the increasing rate of data acquisition. A standard approach for collecting these data is to use dense multi-electrode array (MEA) recordings followed by “spike sorting” algorithms to turn the obtained raw electrical signals into spike trains.

A crucial consideration going forward is the ability to scale to massive datasets–MEAs currently scale up to the order of 10^4^ electrodes, but efforts are underway to increase this number to 10^6^ electrodes^1^. At this scale any manual processing of the obtained data is infeasible. Therefore, automatic spike sorting for dense MEAs has enjoyed significant recent attention [15, 9, 51, 24, 36, 20, 33, 12]. Despite these efforts, spike sorting remains the major computational bottleneck in the scientific pipeline when using dense MEAs, due both to the high computational cost of the algorithms and the human time spent on manual postprocessing.

### Algorithm 1 Pseudocode for the complete proposed pipeline

Input: time-series of electrophysiological data 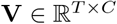, locations 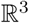

[waveforms, timestamps] ← `Detection`(**V**) % (Section 2.2)

% “Triage” noisy waveforms and collisions (Section 2.4):

[cleanWaveforms, cleanTimestamps] ← `Triage`(waveforms, timestamps)

% Build a set of representative waveforms and summary statistics (Section 2.5)

[representativeWaveforms, sufficientStatistics] ← `coresetConstruction`(cleanWaveforms)

% DP-GMM clustering via divide-and-conquer (Sections 2.6 and 2.7)

[{representativeWaveforms*_i_*, sufficientStatistics*_i_*}*_i_*_=1,…_]

     ← `splitIntoSpatialGroups`(representativeWaveforms, sufficientStatistics, locations)

**for** i=1,… **do** % Run efficient inference for the DP-GMM

    [clusterAssignments*_i_*] ← `SplitMergeDPMM`(representativeWaveforms*_i_*, sufficientStatistics*_i_*)

**end for**

% Merge spatial neighborhoods and similar templates

[allClusterAssignments, templates] ←

    `mergeTemplates`({clusterAssignments*_i_*}*_i_*_=1,…_, {representativeWaveforms*_i_*}*_i_*_=1,…_, locations)

% Pursuit stage to recover collision and noisy waveforms

[finalTimestamps, finalClusterAssignments] ← `deconvolution`(templates)

**return** [finalTimestamps, finalClusterAssignments]

To accelerate progress on this critical scientific problem, our proposed methodology is guided by several main principles. First, *robustness* is critical, since hand-tuning and post-processing is not feasible at scale. Second, *scalability* is key. To feasibly process the oncoming data deluge, we use efficient data summarizations wherever possible and focus computational power on the “hard cases,” using cheap fast methods to handle easy cases. Next, the pipeline should be *modular*. Each stage in the pipeline has many potential feasible solutions, and we improve by rapidly iterating and updating each stage as methodology develops further. Finally, *prior information* is leveraged as much as possible; we share information across neurons and electrodes in order to extract information from the MEA datastream as efficiently as possible.

We will first outline the methodology that forms the core of our pipeline in Section 2.1, and then we demonstrate the improvements in performance on simulated data and a 512-electrode recording in Section 3. We provide additional results in the appendix to further support the details of our methodology.

## 2 Methods

### 2.1 Overview

The inputs to the pipeline are the band-pass filtered voltage recordings from all *C* electrodes and their spatial layout, and the end result of the pipeline is the set of *K* (where *K* is determined by the algorithm) binary neural spike trains, where a “1” reflects a neural action potential from the *k*th neuron. The voltage signals are spatially whitened prior to processing and are modeled as the superposition of action potentials and background Gaussian noise [12]. Succinctly, the pipeline is a multistage procedure as follows: (*i*) detecting waveforms and extracting features, (*ii*) screening outliers and collided waveforms, (*iii*) clustering, and (*iv*) inferring missed and collided spikes. Pseudocode for the flow of the pipeline can be found in Algorithm 1. A brief overview is below, followed by additional details.

Our overall strategy can be considered a hybrid of a matching pursuit approach (similar to that employed by [36]) and a classical clustering approach, generalized and adapted to the large dense MEA setting. Our guiding philosophy is that it is essential to properly handle “collisons” between simultaneous spikes [37, 12], since collisions distort the extracted feature space and hinder clustering, but that matching pursuit methods (or other sparse deconvolution strategies) are relatively computationally expensive compared to clustering primitives. This led us to a “triage-then-cluster-then-pursuit” approach. Our approach “triages” collided or overly noisy waveforms, putting them aside during the feature extraction and clustering stages, and later recovers these spikes during a final “pursuit” or deconvolution stage. The triaging begins during the detection stage in Section 2.2, where we develop a neural network based detection method that significantly improves sensitivity and selectivity. For example, on a simulated 30 electrode dataset with low SNR, the new approach reduces false positives and collisions by 90% for the same rate of true positives. Furthermore, the neural network is significantly better at the *alignment* of signals, which improves the feature space and signal-to-noise power. The detected waveforms then are projected to a feature space and restricted to a local spatial subset of electrodes as in [24] in Section 2.3. Next, in Section 2.4 an outlier detection method further “triages” the detected waveforms and reduces false positives and collisions by an additional 70% while removing only a small percentage of real detections. All of these steps are achievable in nearly linear time. In our simulations, we demonstrate that this large reduction in false positives and collisions dramatically improves accuracy and stability.

Following the above steps, the remaining waveforms are partitioned into distinct neurons via clustering. Our clustering framework is based on the Dirichlet Process Gaussian Mixture Model (DP-GMM) approach [48, 9], and we modify existing inference techniques to improve scalability and performance. Succinctly, each neuron is represented by a distinct Gaussian distribution in the feature space. Directly calculating the clustering on all of the channels and all of the waveforms is computationally infeasible. Instead, the inference first utilizes the spatial locality via *masking* [24] from Section 2.3. Second, the inference procedure operates on a *coreset* of representative points [13] and the resulting approximate sufficient statistics are used in lieu of the full dataset (Section 2.5). Remarkably, we can reduce a dataset with 100*k* points to a coreset of ≃ 10*k* points with trivial accuracy loss. Next, split and merge methods are adapted to efficiently explore the clustering space [21, 24] in Section 2.6. Using these modern scalable inference techniques is crucial for *robustness* because they empirically find much more sensible and accurate optima and permit Bayesian characterization of posterior uncertainty.

For very large arrays, instead of operating on all channels simultaneously, each distinct spatial neighborhood is processed by a separate clustering algorithm that may be run in parallel. This parallelization is crucial for processing very large arrays because it allows greater utilization of computer resources (or multiple machines). It also helps improve the efficacy of the split-merge inference by limiting the search space. This divide-and-conquer approach and the post-processing to stitch the results together is discussed in Section 2.7. The computational time required for the clustering algorithm scales nearly linearly with the number of electrodes *C* and the experiment time.

After the clustering stage is completed, the means of clusters are used as templates and collided and missed spikes are inferred using the deconvolution (or “pursuit” [37]) algorithm from Kilosort [36], which recovers the final set of binary spike trains. We limit this computationally expensive approach only to sections of the data that are not well handled by the rest of the pipeline, and use the faster clustering approach to fill in the well-explained (i.e. easy) sections.

We note finally that when memory is limited compared to the size of the dataset, the preprocessing, spike detection, and final deconvolution steps are performed on temporal minibatches of data; the other stages operate on significantly reduced data representations, so memory management issues typically do not arise here. See Section B.3 for further details on memory management.

### 2.2 Detection

The detection stage extracts temporal windows around action potentials from the noisy raw electrophysiological signal **V** to use as inputs in the following clustering stage. The number of *clean* waveform detections (true positives) should be maximized for a given level of detected collision and noise events (false positives). Because collisions corrupt feature spaces [37, 12] and will simply be recovered during pursuit stage, they are not included as true positives at this stage. In contrast to the plethora of prior work on hand-designed detection rules (detailed in Section C.1), we use a data-driven approach with neural networks to dramatically improve both detection efficacy and alignment quality.

The crux of the data-driven approach is the availability of prior training data. We are targeting the typical case that an experimental lab performs repeated experiments using the same recording setup from day to day. In this setting hand-curated or otherwise validated prior sorts are saved, resulting in abundant training data for a given experimental preparation. In the supplemental material, we discuss the construction of a training set (including data augmentation approaches) in Section C.2, the architecture and training of the network in Section C.3, the detection using the network in Section C.4, empirical performance in Section C.5, and scalability in Section C.5.

A key result is that our neural network dramatically improves the alignment of detected waveforms. This improved alignment improves the fidelity of the feature space and the signal-to-noise power, and the result of the improved feature space can clearly be seen by comparing the detected waveform features from one standard detection approach (`SpikeDetekt` [24]) in Figure 1 (left) to the detected waveform features from our neural network in Figure 1 (middle). Note that the output of the neural net detection is remarkably more Gaussian compared to the output of SpikeDetekt.

**Figure 1:**
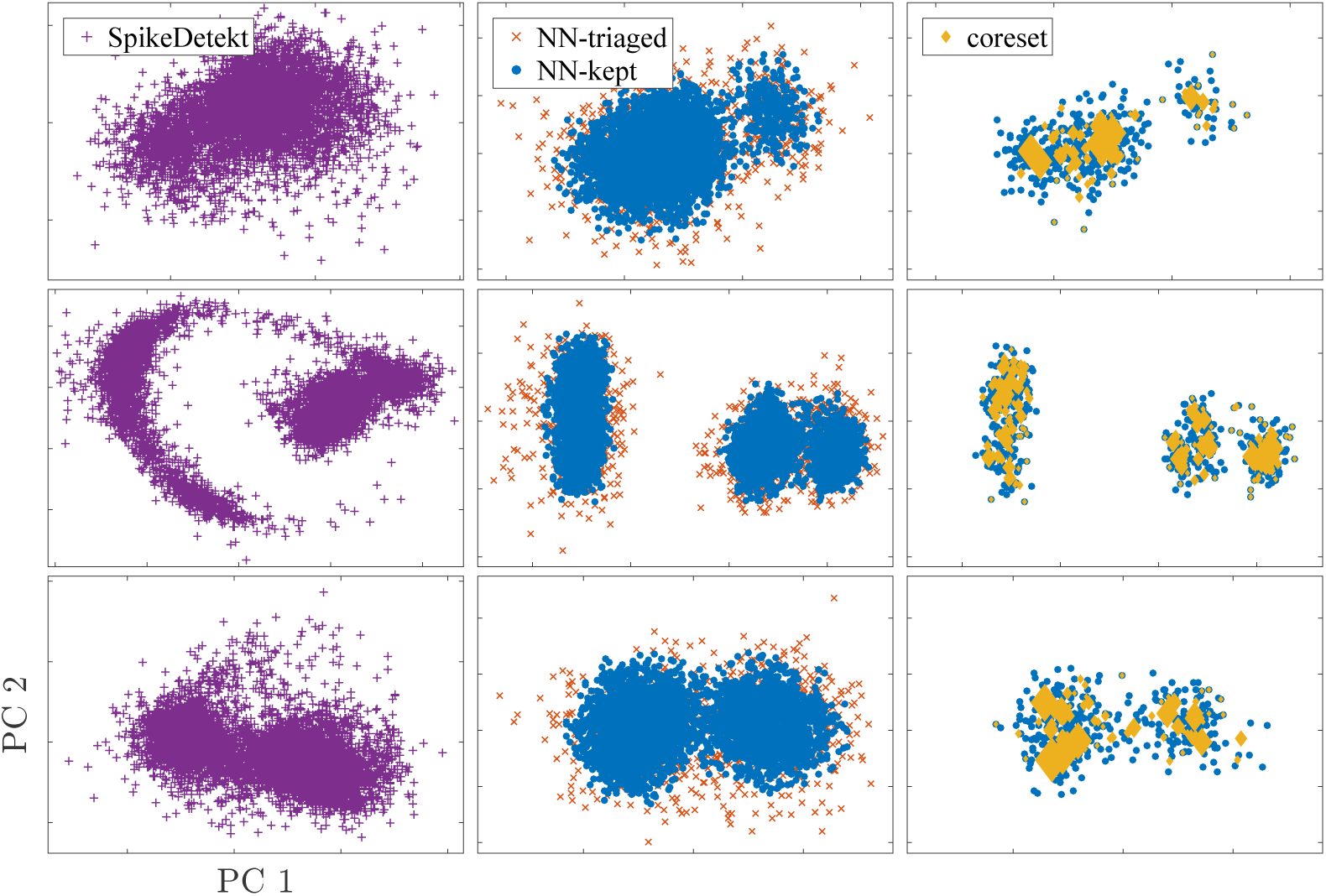
Illustration of Neural Network Detection, Triage, and Coreset from a primate retinal ganglion cell recording. The first column shows spike waveforms from `SpikeDetekt` in their PCA space. Due to poor alignment, clusters have a non-Gaussian shape with many outliers. The second column shows spike waveforms from our proposed neural network detection in the PCA space. After triaging outliers, the clusters have cleaner Gaussian shapes in the recomputed feature space. The last column illustrates the coreset. The size of each coreset diamond represents its weight. For visibility, only 10% of data are plotted.

### 2.3 Feature Extraction and Mask Creation

Following detection we have a collection of *N* events defined as 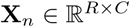 with detection time *t_n_* for *n* = 1, …, *N*. Recall that *C* is the total number of electrodes, and *R* is the number of time samples, in our case chosen to correspond to 1.5ms. Next features are extracted by using uncentered Principal Components Analysis (PCA) on each channel separately with *P* features per channel. Each waveform **X***_n_* is transformed to the feature space **Y***_n_*. To handle duplicate spikes, **Y***_n_* is kept only in its dominant channel, if 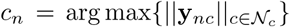, where 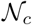 is the local neighborhood of electrodes for the *c^th^* electrode. To address dimensionality increases, spikes are localized by using the sparse masking vector {**m***_n_*} ∊ [0,1]*^C^* method of [24], where the mask should be set to 1 only where the signal exists. The sparse vector reduces the dimensionality and facilitates sparse updates to improve computational efficiency. We give additional mathematical details in Supplemental Section D.

### 2.4 Collision Screening and Outlier Triaging

Many collisions and outliers remain even after our improved detection algorithm. Because these events destabilize the clustering algorithms, the pipeline benefits from a “triage” stage to further screen collisions and noise events. Note that triaging out a small fraction of true positives is a minor concern at this stage because they will be recovered in the final deconvolution step.

We use a two-fold approach to perform this triaging. First, obvious collisions with overlapping spike times and spatial locations are removed. Second, k-Nearest Neighbors (k-NN) is used to detect outliers in the masked feature space based on [27]. To develop a computationally efficient and effective approach, waveforms are grouped based on their primary (highest-energy) channel, and then k-NN is run for each channel. Empirically, these approximations do not suffer in efficacy compared to using the full spatial area. When the dimensionality of *P*, the number of features per channel, is low, a kd-tree can find neighbors in 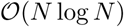 average time. We demonstrate that this method is effective for triaging false positives and collisions in Figure 1 (middle).

### 2.5 Coreset Construction

“Big data” improves density estimates for clustering, but the cost per iteration naively scales with the amount of data. However, often data has some redundant features, and we can take advantage of these redundancies to create more efficient summarizations of the data. Then running the clustering algorithm on the summarized data should scale only with the number of summary points. By choosing representative points (or a “coreset”) carefully we can potentially describe huge datasets accurately with a relatively small number of points [19, 13, 2].

There is a sizable literature on the construction of coresets for clustering problems; however, the number of required representative points to satisfy the theoretical guarantees is infeasible in this problem domain. Instead, we propose a simple approach to build coresets that empirically outperforms existing approaches in our experiments by forcing adequate spatial coverage. We demonstrate in Supplemental Figure S5 that this approach can cover clusters completely missed by existing approaches, and show the chosen representative points on data in Figure 1 (right). This algorithm is based on recursively performing k-means; we provide pseudocode and additional details in in Supplemental Section E. The worst case time complexity is nearly linear with respect to the number of representative points, the number of detected spikes, and the number of channels. The algorithm ends by returning *G* representative points, their sufficient statistics, and masks.

### 2.6 Efficient Inference for the Dirichlet Process Gaussian Mixture Model

For the clustering step we use a Dirichlet Process Gaussian Mixture Model (DP-GMM) formulation, which has been previously used in spike sorting [48, 9], to adaptively choose the number of mixture components (visible neurons). In contrast to these prior approaches, here we adopt a Variational Bayesian split-merge approach to explore the clustering space [21] and to find a more robust and higher-likelihood optimum. We address the high computational cost of this approach with several key innovations. First, following [24], we fit a mixture model on the virtual masked data to exploit the localized nature of the data. Second, following [9, 24], the covariance structure is approximated as a block-diagonal to reduce the parameter space and computation. Finally, we adapted the methodology to work with the representative points (coreset) rather than the raw data, resulting in a highly scalable algorithm. A more complete description of this stage can be found in Supplemental Section F, with pseudocode in Supplemental Algorithm S2.

In terms of computational costs, the dominant cost per iteration in the DPMM algorithm is the calculation of data to cluster assignments, which in our framework will scale at 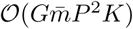, where 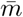 is the average number of channels maintained in the mask for each of the representative points. As a reminder, *G* is the number of representative points and *P* is the number of features per channel. This is in stark contrast to a scaling of 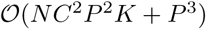 without our above modifications. Both *K* and *G* are expected to scale linearly with the number of electrodes and *sublinearly* with the length of the recording. This leads to a unfortunate dependence on the square of the number of electrodes for each iteration; fortunately, it is feasible to utilize local structure to reduce computations to scale linearly with the number of electrodes. We discuss this approach below in Section 2.7.

### 2.7 Divide and Conquer and Template Merging

Neural action potentials have a finite spatial extent [6]. Therefore, the spikes can be divided into distinct groups based on the geometry of the MEA and the local position of each neuron, and each group is then processed independently. Thus, each group can be processed in parallel, allowing for high data throughput. This is crucial for exploiting parallel computer resources and limited memory structures. Second, the split-and-merge approach in a DP-GMM is greatly hindered when the numbers of clusters is very high [21]. The proposed divide and conquer approach addresses this problem by greatly reducing the number of clusters within each subproblem, allowing the split and merge algorithm to be targeted and effective.

To divide the data based on the spatial location of each spike, the primary channel *c_n_* is determined for every point in the coreset based on the channel with maximum energy, and clustering is applied on each channel. Because neurons may now end up on multiple channels, it is necessary to merge templates from nearby channels as a post-clustering step. When the clustering is completed, the mean of each cluster is taken as a template. Because waveforms are limited to their primary channel, some neurons may have “overclustered” and have a distinct mixture component on distinct channels. Also, overclustering can occur from model mismatch (non-Gaussianity). Therefore, it is necessary to merge waveforms. Template merging is performed based on two criteria, the angle and the amplitude of templates, using the best alignment on all temporal shifts between two templates. The pseudocode to perform this merging is shown in Supplemental Algorithm S3. Additional details can be found in Supplemental Section G.

### 2.8 Recovering Triaged Waveforms and Collisions

After the previous steps, we apply matching pursuit [36] to recover triaged waveforms and collisions. We detail the available choices for this stage in Supplemental Section I.

## 3 Performance Comparison

We evaluate performance to compare several algorithms (detailed in Section 3.1) to our proposed methodology on both synthetic (Section 3.2) and real (Section 3.3) dense MEA recordings. For each synthetic dataset we evaluate the ability to capture ground truth in addition to the per-cluster stability metrics. For the ground truth, inferred clusters are matched with ground truth clusters via the Hungarian algorithm, and then the per-cluster accuracy is calculated as the number of assignments shared between the inferred cluster and the ground truth cluster over the total number of waveforms in the inferred cluster. For the per-cluster stability metric, we use the method from Section 3.3 of [5] with the rate scaling parameter of the Poisson processes set to 0.25. This method evaluates how robust individual clusters are to perturbations of the dataset. In addition, we provide runtime information to empirically evaluate the computational scaling of each approach. The CPU runtime was calculated on a single core of a six-core i7 machine with 32GB of RAM. GPU runtime is given from a Nvidia Titan X within the same machine.

### 3.1 Competing Algorithms

We compare our proposed pipeline to three recently proposed approaches for dense MEA spike sorting: KiloSort [36], Spyking Circus [51], and MountainSort [31]. Kilosort, Spyking Cricus, and MountainSort were downloaded on January 30, 2017, May 26th, 2017, and June 7th, 2017, respectively. We dub our algorithm Yet Another Spike Sorter (`YASS`). We discuss additional details on the relationships between these approaches and our pipeline in Supplemental Section I. All results are shown with no manual post-processing.

### 3.2 Synthetic Datasets

First, we used the biophysics-based spike activity generator `ViSAPy` [18] to generate multiple 30-channel datasets with different noise levels and collision rates. The detection network was trained on the ground truth from a low signal-to-noise level recording. Then, the trained neural network is applied to all signal-to-noise levels. The neural network dramatically outperforms existing detection methodologies on these datasets. For a given level of true positives, the number of false positives can be reduced by an order of magnitude. The properties of the learned network are shown in Supplemental Figure S3 and the ROC curves are shown in Supplemental Figure S4.

Performance is evaluated on the known ground truth. For each level of accuracy, the number of clusters that pass that threshold is calculated to demonstrate the relative quality of the competing algorithms on this dataset. Empirically, our pipeline (`YASS`) outperforms other methods. This is especially true in low SNR settings, as shown in Figure 2. The per-cluster stability metric is also shown in Figure 2. The stability result demonstrates that `YASS` has significantly fewer low-quality clusters than competing methods.

**Figure 2:**
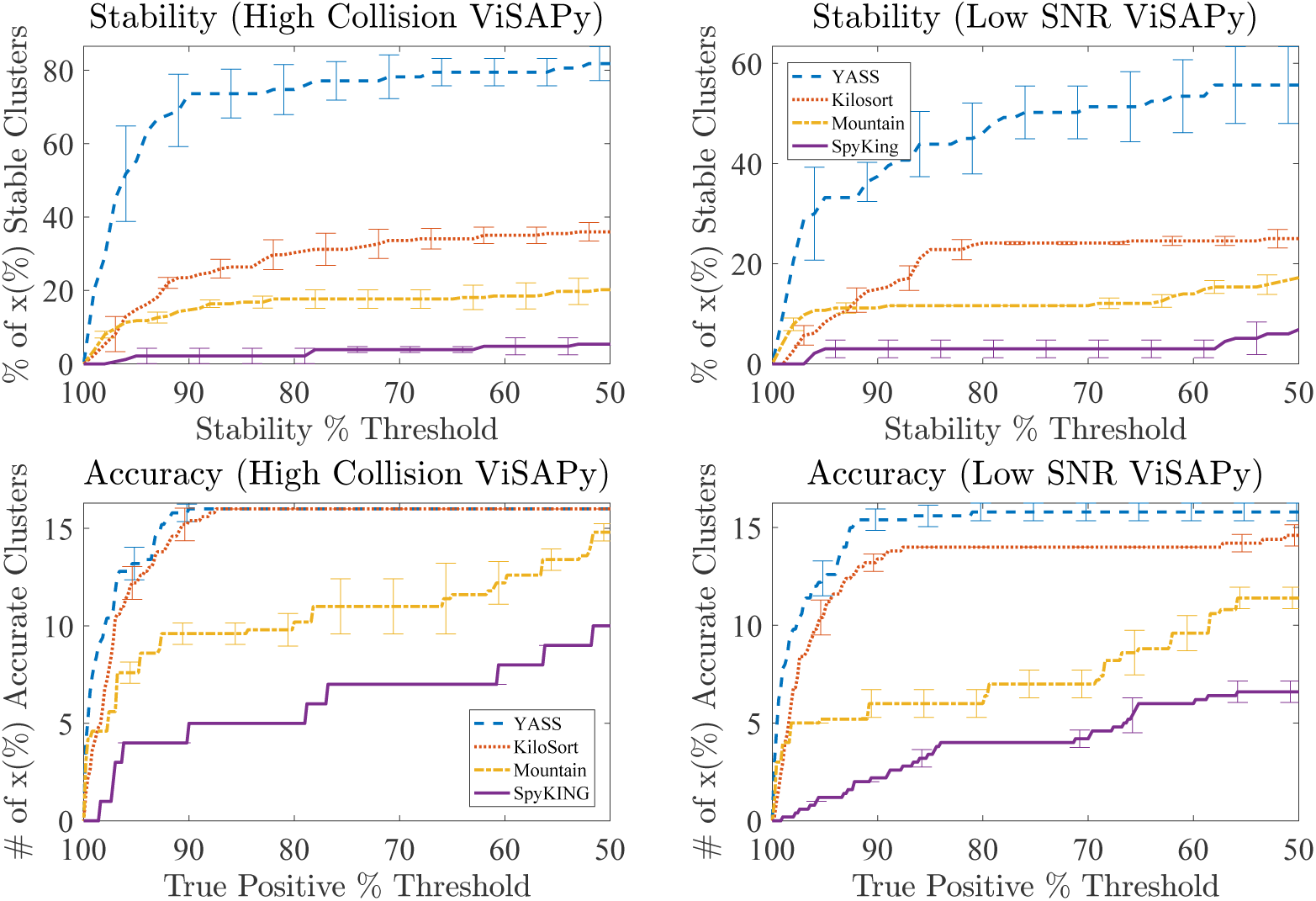
Simulation results on 30-channel ViSAPy datasets. Left panels show the result on `ViSAPy` with high collision rate and Right panels show the result on `ViSAPy` with low SNR setting. (Top) stability metric (following [5]) and percentage of total discovered clusters above a certain stability measure. The noticeable gap between stability of `YASS` and the other methods results from a combination of high number of stable clusters and lower number of total clusters. (Bottom) These results show the number of clusters (out of a ground truth of 16 units) above a varying quality threshold for each pipeline. For each level of accuracy, the number of clusters that pass that threshold is calculated to demonstrate the relative quality of the competing algorithms on this dataset. Empirically, our pipeline (`YASS`) outperforms other methods.

### 3.3 Real Datasets

To examine real data, we focused on 30 minutes of extracellular recordings of the peripheral primate retina, obtained ex-vivo using a high-density 512-channel recording array [30]. The half-hour recording was taken while the retina was stimulated with spatiotemporal white noise. A “gold standard” sort was constructed for this dataset by extensive hand validation of automated techniques, as detailed in Supplemental Section H. Nonstationarity effects (time-evolution of waveform shapes) were found to be minimal in this recording (data not shown).

We evaluate the performance of `YASS` and competing algorithms using 4 distinct sets of 49 spatially contiguous electrodes. Note that the gold standard sort here uses the information from the full 512-electrode array, while we examine the more difficult problem of sorting the 49-electrode data; we have less information about the cells near the edges of this 49-electrode subset, allowing us to quantify the performance of the algorithms over a range of effective SNR levels. By comparing the inferred results to the gold standard, the cluster-specific true positives are determined in addition to the stability metric. The results are shown in Figure 3 for one of the four sets of electrodes, and the remaining three sets are shown in Supplemental Section B.1. As in the simulated data, compared to KiloSort, which had the second-best overall performance on this dataset, `YASS` has dramatically fewer low-stability clusters.

**Figure 3:**
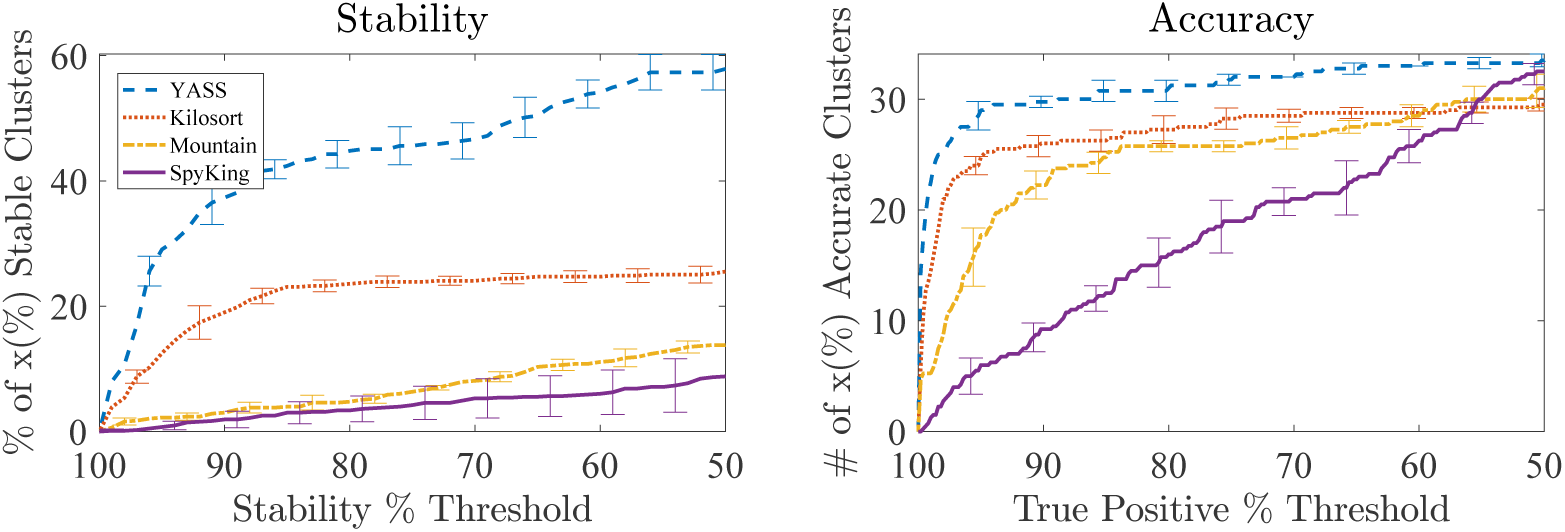
Performance comparison of spike sorting pipelines on primate retina data. (Left) The same type of plot as in the top panels of Figure 2. (Right) The same type of plot as in the bottom panels of Figure 2 compared to the “gold standard” sort. Similar trends

Finally, we evaluate the time required for each step in the `YASS` pipeline (Table 1). Importantly, we found that `YASS` is highly robust to data limitations: as shown in Supplemental Figure S2 and Section B.2, using only a fraction of the 30 minute dataset has only a minor impact on performance. We exploit this to speed up the pipeline. Remarkably, running primarily on a single CPU core (only the detect step utilizes a GPU here), `YASS` achieves a several-fold speedup in template and cluster estimation compared to the next fastest competitor^2^, Kilosort, which was run in full GPU mode and spent about 30 minutes on this dataset. We plan to further parallelize and GPU-ize the remaining steps in our pipeline next, and expect to achieve significant further speedups.

**Table 1:** Running times of the main processes on 512-channel primate retinal recording of **30 minutes duration.** Results shown using a single CPU core, except for the detection step (2.2), which was run on GPU. We found that full accuracy was achieved after processing just one-fifth of this dataset, leading to significant speed gains. Data Extraction refers to waveform extraction and Performing PCA (2.3). Triage, Coreset, and Clustering refer to 2.4, 2.5, and 2.6, respectively. Template Extraction describes revisiting the recording to estimate templates and merging them (2.7). Each step scales approximately linearly (Section B.2).

| Detection (GPU) | Data Ext. | Triage | Coreset | Clustering | Template Ext. | Total |
| --- | --- | --- | --- | --- | --- | --- |
| 1m7s | 42s | 11s | 34s | 3m12s | 54s | 6m40s |

## 4 Conclusion

`YASS` has demonstrated state-of-the-art performance in accuracy, stability, and computational efficiency; we believe the tools presented here will have a major practical and scientific impact in large-scale neuroscience. In our future work, we plan to continue iteratively updating our modular pipeline to better handle template drift, refractory violations, and improved strategies for collision deconvolution.

## Acknowledgements

This work was partially supported by NSF grants IIS-1546296 and NSF IIS-1430239.

## A Notation

The following notation is employed: scalars are lowercase italicized letters, e.g. *x*, constants such as max indices are represented by uppercase italicized letters, e.g. *N*, vectors are bolded lowercase letters, e.g. x, and matrices are bolded uppercase letters, e.g. **X**. Major notations used in the paper are summarized in Table S1.

**Table S1:** Summary table of notation used within the manuscript.

| Notation | Explanation | Default value (if exists) |
| --- | --- | --- |
| **Data Constants** |  |  |
| $T$ | Recording length | |
| $C$ | Total number of channels | |
| $N$ | Number of detected spikes | |
| $t_n$ | Temporal location of spike $n$ | |
| $c_n$ | Index of the main channel of spike $n$ | |
| $C_{eff}$ | Number of neighboring channels | |
| $G$ | Number of representative points after the Coreset algorithm | |
| **Data structures** |  |  |
| $\mathbf{V} \in \mathbb{R}^{T \times C}$ | Recording | |
| $\mathbf{X}_n \in \mathbb{R}^{R \times C}$ | Voltage trace (waveform) of spike $n$ | |
| $\mathbf{m}_n \in [0, 1]^C$ | Masking vector for spike $n$ | |
| $\mathbf{Y}_n \in \mathbb{R}^{P \times C}$ | $\mathbf{X}_n$ mapped to the feature (PCA) space | |
| $\mathbf{x}_{nc}, \mathbf{y}_{nc}$ | $c^{\text{th}}$ column of $\mathbf{X}_n, \mathbf{Y}_n$ | |
| $m_{nc}$ | $c^{\text{th}}$ entry of $\mathbf{m}_n$ | |
| $\tilde{\mathbf{X}}_n, \tilde{\mathbf{x}}_{nc}, \tilde{\mathbf{Y}}_n, \tilde{\mathbf{y}}_{nc}$ | Virtual data distribution of $\mathbf{X}_n, \mathbf{x}_{nc}, \mathbf{Y}_n, \mathbf{y}_{nc}$ | |
| $\mathbf{P} \in \mathbb{R}^{T \times C}$ | Probability output of NN | |
| $\mathbf{W}_k \in \mathbb{R}^{R \times C}$ | Template of cluster $k$ | |
| $\mathbf{m}_k^w \in \{0, 1\}^C$ | Mask of cluster $k$ | |
| **Parameters** |  |  |
| $R$ | Temporal window size of spike | 1.5 ms |
| $R'$ | Half of window size | |
| $P$ | Per channel dimension of data in PCA domain | 3 |
| $\tau$ | probability threshold on $\mathbf{P}$ | 0.5 |
| $\theta_w, \theta_s$ | weak and strong threshold for masking | $F_{\chi^2, P}^{-1}(0.5), F_{\chi^2, P}^{-1}(0.9)$ |
| $F_{\chi^2, df}^{-1}$ | Inverse CDF of Chi-squared Distribution with $df$ d.f. | |
| $K_{++}$ | Number of Clusters in Kmeans++ for Coreset | 10 |
| $D_{max}$ | Distance threshold for Coreset | $2F_{\chi^2, PC_{eff}}^{-1}(0.9)$ |
| $m_0, \lambda_0, \mathbf{W}_0, v_0$ | Prior parameters for the DP-GMM Normal-Whishart | $\mathbf{0}, 0.01, \frac{1}{v_0} \mathbf{I}_P, P + 2$ |
| $\alpha_0$ | Prior parameter for beta dist. in stick-breaking in DP-GMM | 1 |
| $s_{max}$ | Maximum shift allowance for template merging | 0.5 ms |
| $\theta_1$ | threshold on cosine angle in template merging | 0.85 |
| $\theta_2$ | threshold on size in template merging | 0.6 |

## B Additional results

In this section, we first provide the performance metrics on the other three sets of 49-channel recordings of the primate retina referenced from Section 3.3. Next, we describe how limiting the temporal duration of the data effects performance.

### B.1 Results on Three Additional Real 49-Electrode Sets of Data

The summary performance metrics for the three additional recordings of the primate retina are reported in Figure S1. Note that `YASS` outperforms the other methods, especially in terms of stability of the clusters. This is due to `YASS` providing both more stable clusters and fewer total clusters in general.

**Figure S1:**
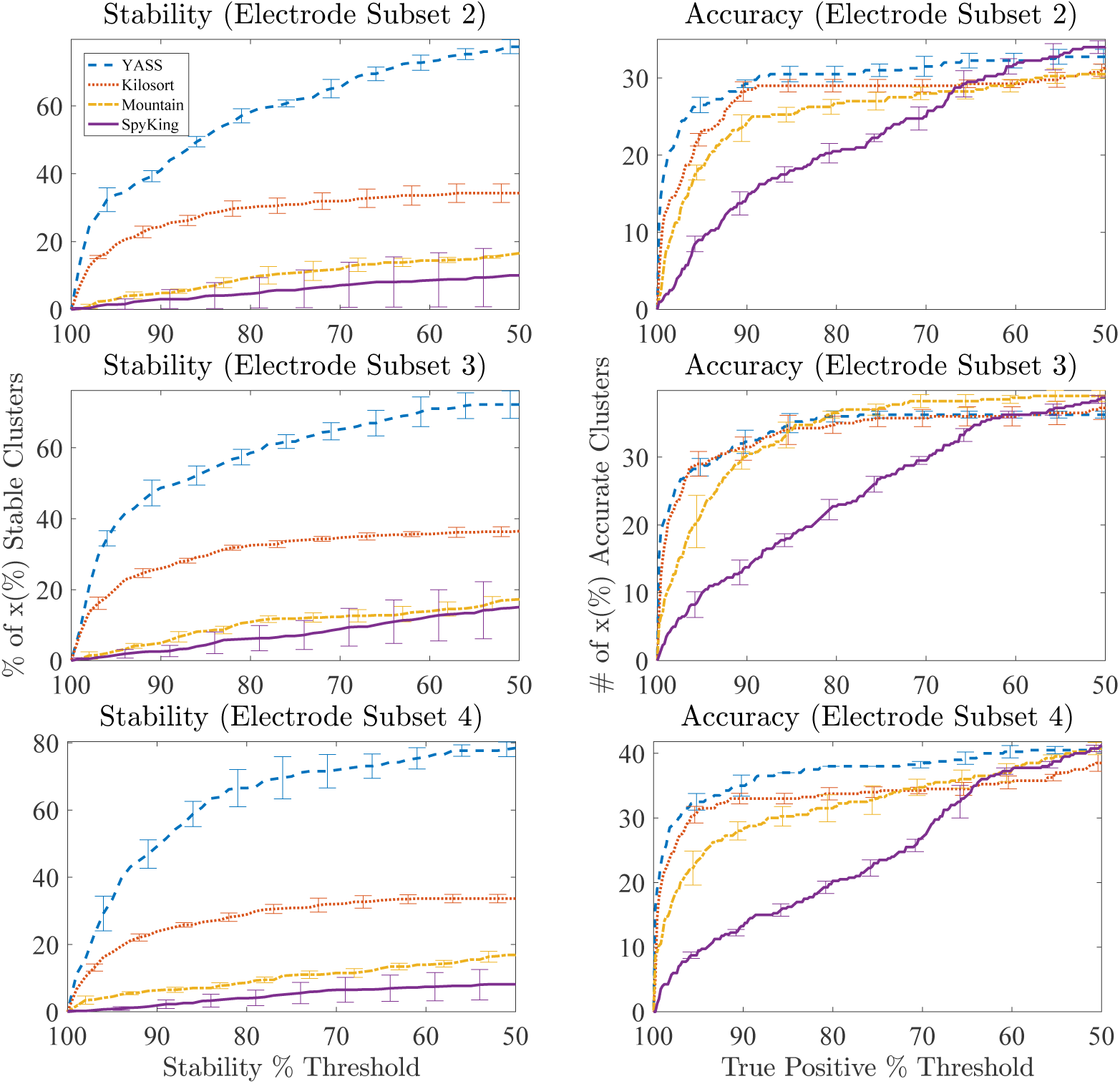
Comparison of performance on the other three 49 channel datasets from primate retina. Each row corresponds to one of the three additional datasets. Figures on the left depict stability metric and what percentage of total discovered clusters are above the chosen stability threshold on the x axis. Figures on the right depict the true positive accuracies with respect to partial ground truth and how many discovered clusters are above the chosen true positive accuracy.

### B.2 Accuracy with Respect to Data Length

The effect of using a partial recording to estimate waveform templates is investigated further. The summary is described in the left and right panels of Figure S2. To illustrate the effect, the full recording is randomly subsetted by the specified length. Waveform templates are estimated using information only from the subset. Accuracy loss becomes insignificant with more than 20% of the full length on a 30 minute dataset.

**Figure S2:**
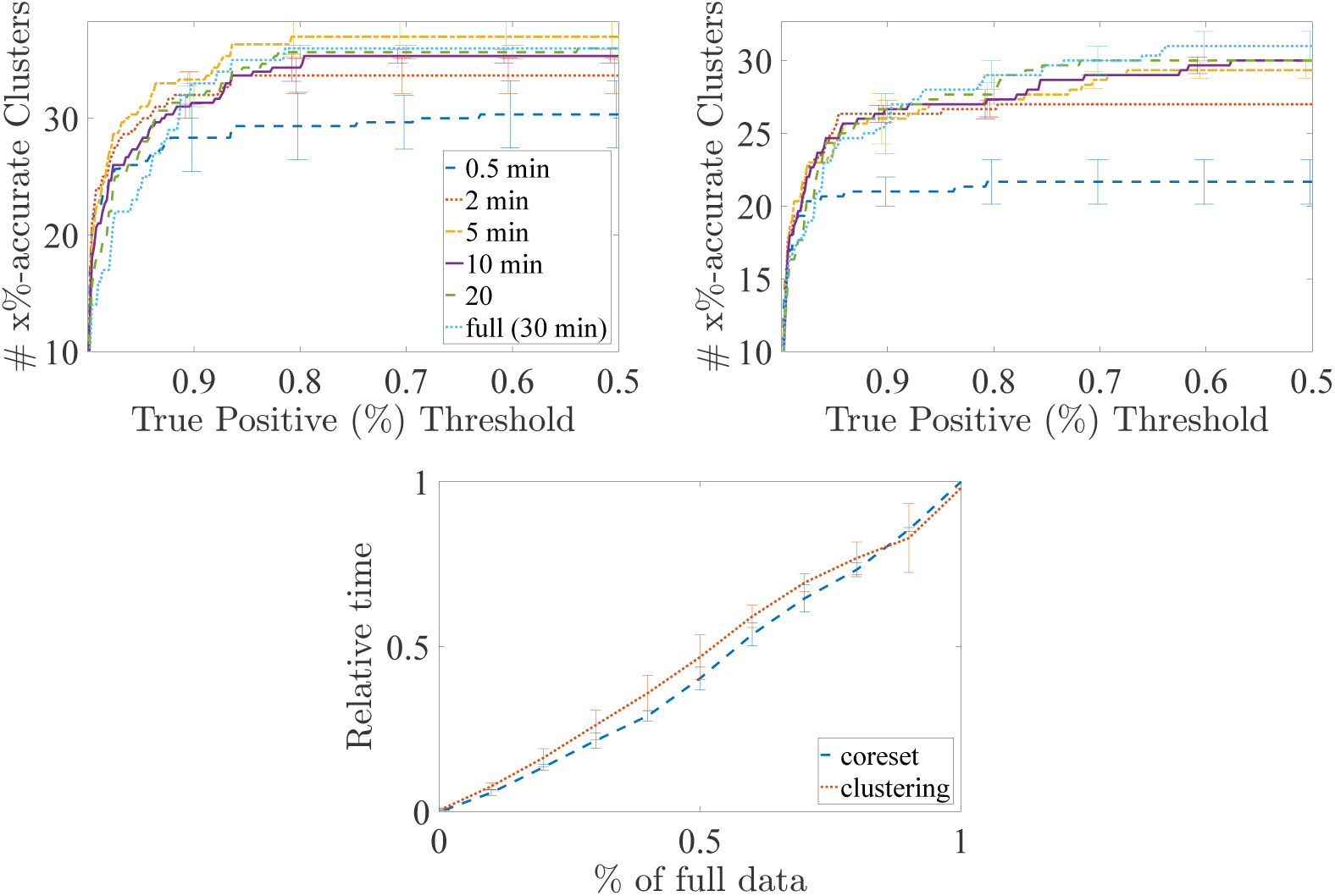
Using only a portion of data to estimate templates. (Top) `YASS` is tested on two sets of recording with 49 channels and 30 minutes length from the retinal cells. Only a portion of data is randomly extracted and templates are estimated. As shown, extracting only 5-10 minutes was enough to produce similar performance as on the full dataset. (Bottom) The scalability of `YASS` is shown. As shown, the coreset and clustering algorithms are roughly linear in computational costs with increasing time.

Furthermore, the scalability of `YASS` is demonstrated in the bottom panel of Figure S2. As illustrated, the main parts of the full pipeline, coreset and clustering, scale almost linearly.

### B.3 When Data Exceeds Memory

Large recordings exceed the memory capacity of typical workstations. This issue in handled in the preprocessing and spike detection by temporally partitioning the recording and processing each temporal subsection individually. Afterwards, the divide and conquer approach from Section 2.7 significantly reduces the memory requirement by reducing the number of waveforms and their spatial extent. If the memory limits are still exceeded due to extremely long recordings, data are randomly subsetted and postprocessed.

Another possible approach is that, following the result of Section B.2, data can be processed partially up to the point where it does not exceed memory. Once the templates are estimated based on the partial recording, deconvolution should handle spikes from the remainder of recording. We have not yet needed to implement this approach, however.

## C Additional details on the Detection Algorithm

### C.1 Relationships to Existing Detection Algorithms

The goal of a detection algorithm within a spike sorting pipeline is to extract (unsorted) action potentials from the raw electrophysiological signal to use as inputs for a downstream clustering algorithms. It is crucial for the subsequent steps of the pipeline that the detected action potentials cover all present neural shapes with few false positives, where false positives here are defined as either noise events, collisions (two or more waveforms simultaneously occurring in time or space), or poorly aligned spikes. Historically, most research labs have used a simple voltage threshold to determine whether a section of signal should be considered an action potential [29], but many other decision rules have been considered, such as the nonlinear energy operator [32] and wavelet thresholding [50].

Most proposed detection rules above operated on a single channel at a time (although Bayesian optimal detection has been used on multiple channels [8, 33]). A simple approach that works either for a single channel or for many channels is simply to use template matching [4]. However, template matching requires having templates that are specific to the recording of interest in advance and does not allow much variability in spike shape. Even in a repeated experiment setting, small changes in the environment, such as shift in electrodes, would change the shape of templates and, thus, render templates obtained from a previous experiment unusable. This viewpoint is taken in several approaches, such as [28, 12, 36]. Despite the appeal of these approaches, they are often computationally expensive and difficult to combine with state-of-the-art clustering approaches.

With the increasing popularity of dense MEAs, more complex rules have been proposed to utilize information from all channels simultaneously, such as `SpikeDetekt` [24]. We note that our methodology is structured in a modular way, such that our pipeline can easily adopt any of these existing methods. However, we also advocate for the development of *data-driven* approaches. In many cases the same device types have been reused for many experiments, and there exists a large collection of example data where false positives and true positives have been thoroughly assessed and curated. In these cases, we will propose a novel approach based on recent advances in deep learning to learn efficient, real-time detection algorithms. This is in contrast to many existing approaches where features hand-tuned (e.g. threshold, NEO, `SpikeDetekt`).

There have been some previous data driven efforts to train a detection algorithm. For example, [25] used hand-curated results from previous sorts to train a neural network, but this was used to classify waveforms rather than as a pure detection method. Furthermore, [45] trained a support vector machine to detect spikes in a simulated recording that provided improvements over a threshold method. Our approach is down a similar line, where we use both previous hand-curated results and synthetic data to train a neural network that dramatically improves the detection quality, as demonstrated in Figure S4 compared to `SpikeDetekt`.

**Figure S3:**
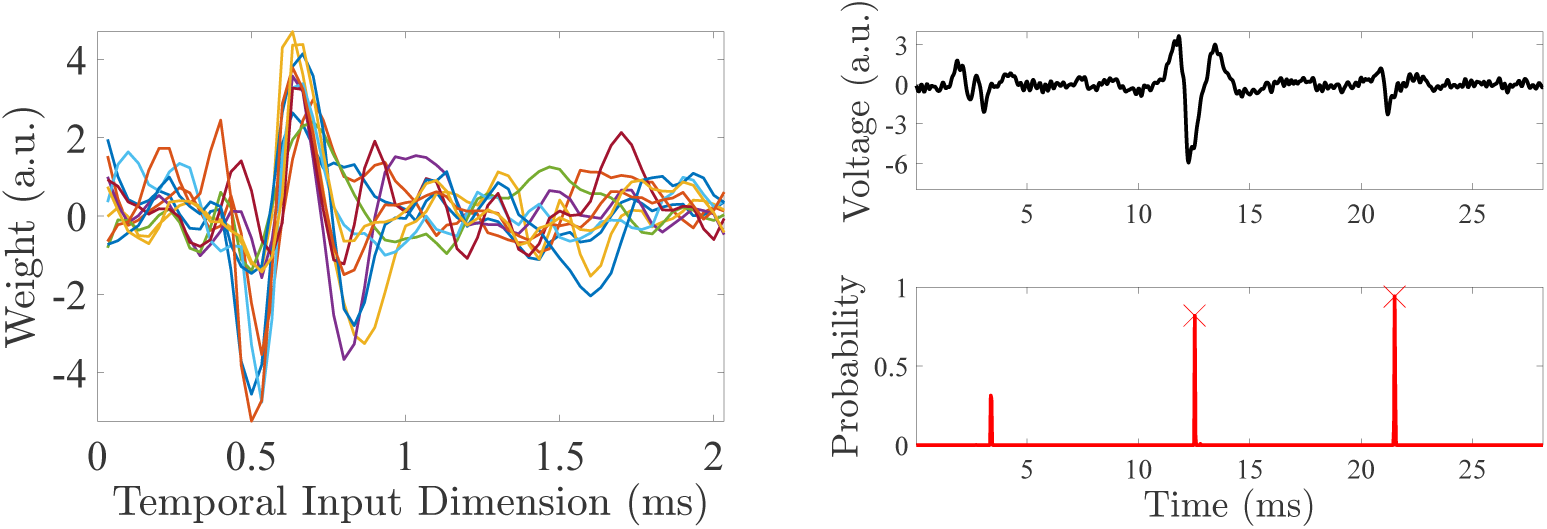
Illustration of the Neural Network Detection. (Left) The 10 learned filters in the first convolutional layer of the neural network. (Right) The neural network transforms a neural recording (Top) into probabilities of spikes (Bottom). Locations of isolated spikes clearly have high probabilities.

**Figure S4:**
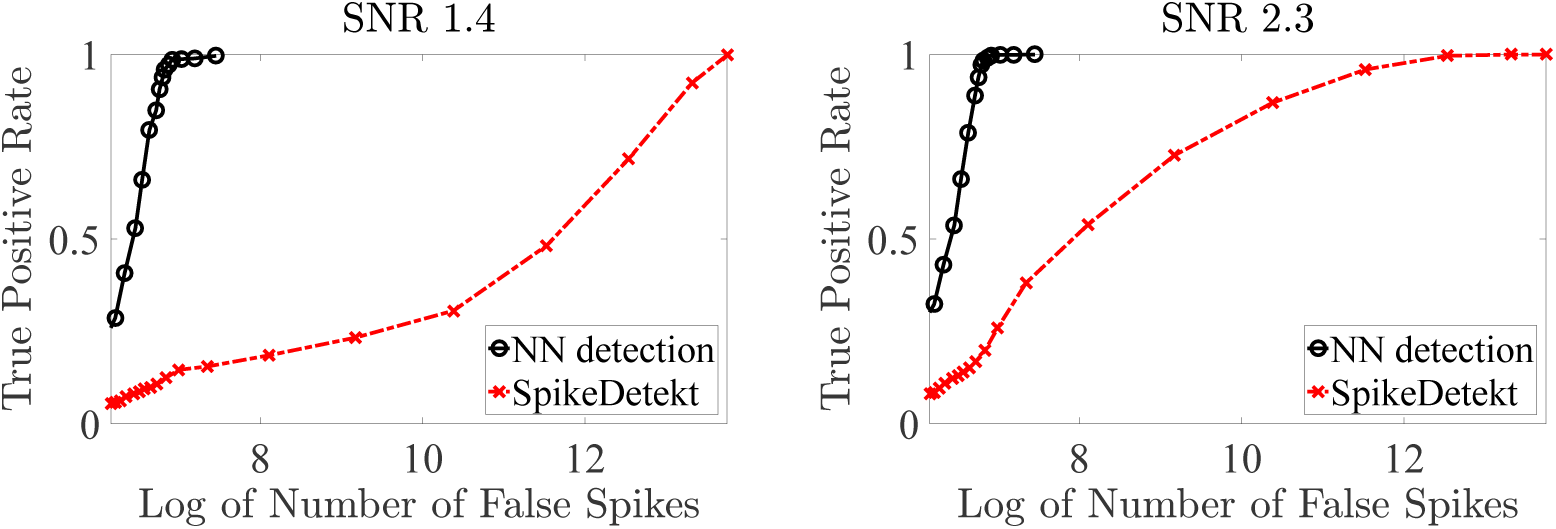
Receiver Operating Characteristic (ROC) curve for Neural Network Detection and SpikeDetekt. The left panel shows the ROC curve for a cluster with SNR 1.4 and the right panel shows for a cluster with SNR 2.3. Note that instead of the false positive rate, the number of false positive spikes are used for comparison.

While this approach is dependent on existing training data and may not be practical everywhere, we emphasize that our pipeline gives state-of-the-art or near state-of-the-art results conditioned on the spike detection method. When curated training data exists (which is true for many research labs), though, this approach will learn features necessary for detection from the data, and we demonstrate that it can *significantly* improve performance in real data problems. It dramatically reduces the amount of false positives for the same level of true spikes. More importantly, by detecting only well isolated spikes and aligning them properly, it improves the quality of the feature extraction and the signal-to-noise ratio.

### C.2 Neural Network Training Data

The training data for the neural network is constructed from previous sorts. Training labels are either defined from a full deconvolution pipeline or a hand-curated effort to validate results. We first focus on a single channel to facilitate training and improve generalization. Specifically, we assume that we have *N_train_* time series, where 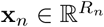 reflects the noisy voltage signal and **y** ∊ {0, 1}*^Rn^* is a binary time series where the value “1” denotes the presence of a well-isolated action potential (i.e. no collisions). We assume that *R_n_ >> R*, where *R* is the length of an action potential.

It is likely that previous sorts do not reflect a perfect ground truth. They may contain false negatives and poorly aligned spikes, which could lead to the creation of faulty training set. As an alternative, synthetic spikes can be constructed from the previous sorts [49, 43].

We also provide a simple method to augment the training set which requires that the mean of clusters must resemble the shape of spikes. A “noise” training set can be obtained from the noise floor, which is determined by using a low amplitude threshold so that it excludes most of spikes. As a low threshold is used, it is rescaled to match the real noise level presented in the recording. An augmented spike train is constructed by superimposing real noise onto randomly-scaled templates. This way, spikes are well aligned and also vary in size.

### C.3 Neural Network Structure

The architecture of the detection network is a fully convolutional neural network with two hidden layers (see [17] for a background on neural networks). There are *K*_1_, *K*_2_, and *K*_3_ filters of length *R*_1_, *R*_2_, and *R*_3_, respectively, in each of the convolutional filter banks. In our experiment, *R*_1_ = *R*, which corresponds to 2 milliseconds (*K*_1_ = 60 for 30kHz recording) and *R*_2_, *R*_3_ = 5. *K*_1_, *K*_2_, and *K*_3_ are set to 10, 5, and 1 respectively. The input at the first layer is the electrophysiological time series on a single channel, and the output is the binary labels. A rectified linear unit nonlinearity, defined as *ReLU*(*x*) = max(*x*, 0), is used at each hidden layer. A sigmoid nonlinearity *σ*(*x*) = exp(*x*)*/*(1 + exp(*x*)) was used at the output to map the probability to the [0, 1] space. The model was regularized by adding an *ℓ*_2_ penalty on the filter weights. The parameters are learned using the Adam algorithm [26] with the parameter settings to minimize the cross-entropy training loss.

### C.4 Detection using the Neural Network

After the neural network has been learned, it is applied in a channel-wise manner to transform the recorded voltages 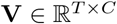 into a matrix of probabilities **P** ∊ [0, 1]*^T^*^×*C*^. A spike is declared if the maximum value of **P** passes a threshold *τ* in a local temporal area; spatial overlaps are handled in a following step as discussed in Section 2.3. The threshold *τ* is tunable to alter tradeoff between false positives and true positives. In our experiments, *τ* is simply set to 0.5.

To capture the temporal windows, each waveform includes 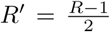 samples before and after the spike time (corresponding to 1 milliseconds). For each *n* of the *N* detections, let *t_n_* and *c_n_* be temporal location and spatial location of the waveform. Each spike is then defined as 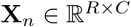, where the *c^th^* channel is defined 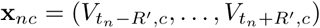.

### C.5 Empirical Performance and Scalability

The performance of the proposed detection algorithm is empirically tested and compared to `SpikeDetekt` on simulated data with high noise level using `ViSAPy`. Since spikes with high enough energy are captured well, cluster-specific ROC curves are plotted. As shown in Figure S4, using the neural network for detection dramatically reduces the number of false positives while keeping the number of true positives reasonably high. Here, SNR is defined as 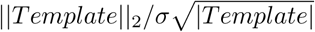, where *Template* is the mean of all waveforms in the cluster at its highest energy channel and *σ* is the noise standard deviation.

When the trained neural network is applied, the algorithm scales linearly with the time duration of the sample and the number of channels. As an example of the computational needs of applying this step, on a 5 minute recording of 49 channels, the proposed detection algorithm requires 6 seconds on a single NVIDIA Titan X GPU. Training the network is more time consuming, but not a computational bottleneck. For a 512 channel recording used in our experiments (described in Section 3), the training process required 2 minutes on the same GPU. Compared to the total spike sorting time, as discussed in Section 3.3, training the network is a reasonable cost that quickly becomes trivial as the algorithm is applied to more and more datasets.

## D Additional Details on Feature Extraction and Masking

The data dimensionality of each spike increases linearly with the number of channels, which naively poses huge computational challenges and renders real-time analysis of large MEAs infeasible. Therefore, it is necessary to utilize spatial *locality*: a given neuron will appear strongly on only a subset of neighboring electrodes. This spatial locality is constructed by restricting detected waveforms to nearby channels via a sparse masking vector, {**m***_n_*} = [**m***_n_*_1_*, …*, **m_nC_**] ∊ [0, 1]*^C^*, as in [24]. This masking vector should be 1 on channels where the neural waveform is reasonably strong (i.e. only the local area), and 0 otherwise. Once the sparse representation is used, the waveform only needs to be considered on channels where the mask is 1, dramatically reducing the effective dimensionality. The effective dimensionality is then dependent on the spatial density of the array and not its total size, so increasing the size of the array does not alter the effective dimensionality. This is a crucial consideration for computational efficiency in many steps of the pipeline, where there is a linear (or worse) scaling with the effective dimensionality.

The construction of the sparse masking vector follows from [24]. First, note that in the feature space a channel that does not have waveform signal present (i.e. a “noise” channel) is expected to follow the normal distribution 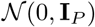. This distribution follows from the assumption of whitened background noise. Therefore, the task is to decide whether the signal on each channel is simply a noise event or not. Using a “strong” and “weak” threshold, *θ_s_* and *θ_w_*, respectively, each mask entry for waveform *n* in each channel *c* is determined based on the norm of the power in the feature space, given by

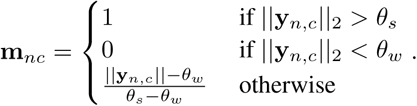

Note that there is additional spatial connectivity considered in [24], which is ignored here. In our empirical results, there was no additional benefit to considering spatial connectivity and it added significant extra computational time.

To facilitate efficient inference, the detected waveforms are represented by a “virtual” data distribution, where 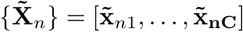, which is given by

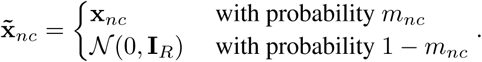

Thus, virtual data are given by a mixture of either the original value or a draw from the noise distribution. *R* is the temporal length of each waveform. Remarkably, the virtual data distribution allows trivial updates during the clustering stage whenever *m_nc_* is 0 [24]. This is crucial to make the computational costs in the DP-GMM scale linearly with the number of electrodes and the length of the recording, as discussed in Section 2.6. The same approach is used to construct a virtual data distribution in the feature space, which would be used in the clustering algorithm.

## E Additional details on the Coreset Construction

There is significant prior work on how to develop a set of representative points that make up a coreset [19, 13]. These prior works developed rigorous approaches with statistical guarantees on representation fidelity. After the development of the coreset, the clustering algorithm is then run on the summary data in time that scales with the size of the coreset, *G* rather than the raw data size. However, in practice, we found the resulting *G* to not be large enough to provide reasonable guarantees. Furthermore, following existing strategies with a smaller number of representative points empirically failed to provide coverage over all clusters, as shown in Figure S5.

**Figure S5:**
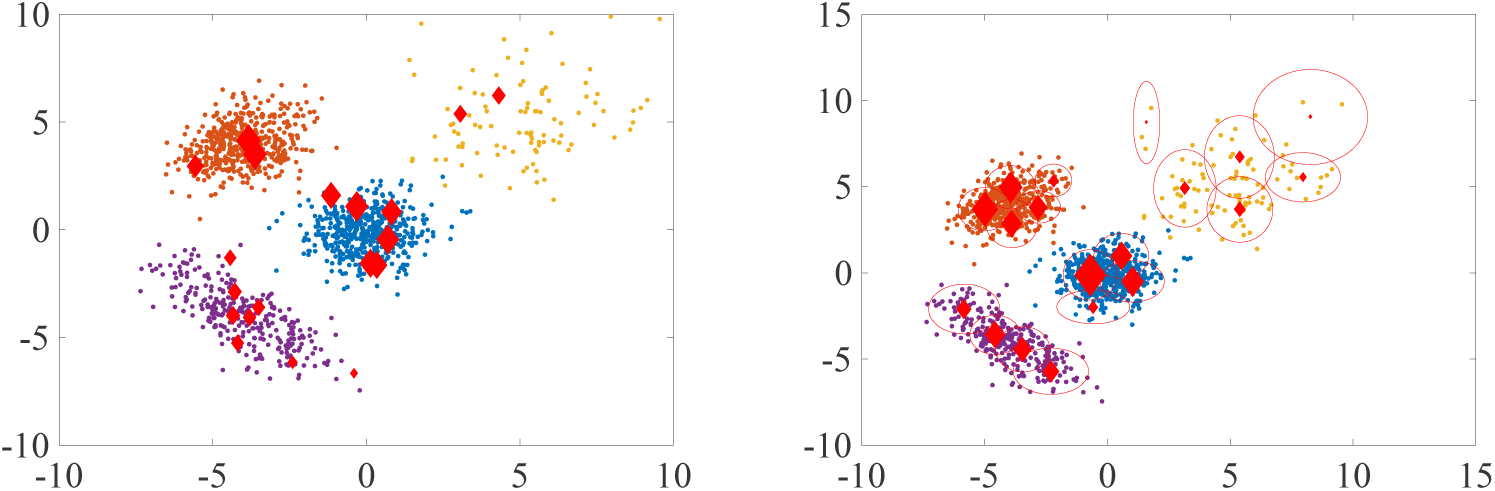
Illustration of coreset construction. Simulated data from Gaussian Mixture models. Four different clusters are clearly visible. (Left) Coreset from [2]. The red points are the coreset and their size represents the weight. It is clear that the cluster shapes will not be well represented by the existing coreset method. (Right) Coreset from the proposed method. The sizes of red diamonds represent the weights and the circles are two times standard deviations of each group. As all points contribute to the coreset, the shape of cluster will be well preserved.

Thus we provide a simple alternative method to construct the coreset that empirically worked well to capture all visible clusters. The procedure begins by first running K-means++ [1] based on the Euclidean distance with a predefined number of clusters, *K*_++_. The running time of this step is 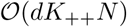 if we assume constant iterations, as is common [1, 3]. *d* here represents the complete dimensionality of each point, which is *CP* without considering the effect of the masking vector. To finish the construction of the coreset, if any center in k-means has a associated point that is unacceptably far away (determined by a threshold *D*_max_), each cluster is recursively partitioned by reapplying K-means++. At the end, only sufficient statistics, mean and covariance of each partition, need to be passed on to DP-GMM described in the next section. The details of this recursive strategy are shown in Algorithm S1.

Unfortunately, this approach is infeasible to run on all channels simultaneously. To address this problem, data are partitioned based on their primary channels (the channel with the highest energy), and only the waveform data on the primary channel and its neighbors are used to construct the coreset. Algorithm S1 is then applied on each set (primary channel) in the partition. These approach reduces the complexity of the primary K-means++ call to 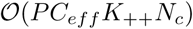, where *C_e f f_* is the number of neighboring channels and *N_c_* is the number of waveforms in the *c*th partition. Note that *K*_++_ is set to a smaller value when applied to a single channel, providing a large source of computational savings.

### Algorithm S1 Constructing the Coreset of Representative Points

[representativeWaveforms, sufficientStatistics] ← `coresetConstruction`(cleanWaveforms)

Input: cleanWaveforms are given by 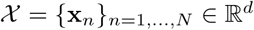

Algorithmic Settings: Distance threshold *D*_max_, number of splits *K*, and the maximum

    iteration number *I_max_*, distance function *D*(·, ·)

Output: Representative waveforms and their sufficient statistics

Apply `coreset` (below) to partition the data

Return: centroids (representativeWaveforms) and sufficient statistics of each entry in the partition

Support function: 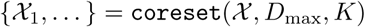

% Run initial partition

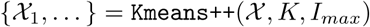

% Recursively split partitions that are too diffuse

**for** *k* = 1, …, *K* **do**

    **if** 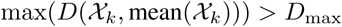 **then**

         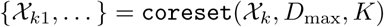

    **end if**

**end for**

Gather all partitions and reconstruct into 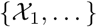

**for** *k* = 1, … **do**

    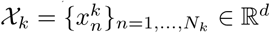

    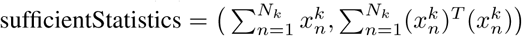

**end for**

## F Additional details on the Dirichlet process Gaussian mixture model (DP-GMM)

One of the biggest issues with using GMMs for clustering is that the appropriate number of mixture components *k* (the number of visible distinct neural signals) is unknown *a priori*. A common approach is to fit a GMM with varying values of *k* and then perform model selection through the Akaike Information Criterion or the Bayesian Information Criterion [24, 44].

A contrasting approach is the Dirichlet Process Gaussian Mixture Model (DP-GMM), which defines a prior over *k*. There is a rich literature on inferring *k* using either Markov chain Monte Carlo methods or variational inference; see [48, 9] for previous spike sorting applications. Here, we will first set up the DP-GMM and describe a split-merge Variational inference method to learn the model based on [21]. We will then describe how to alter these algorithms to work with our coreset to facilitate fast and scalable inference. All GMM approaches require inference over a non-convex log-likelihood, where finding optimal parameters is non-trivial. The split-merge approach empirically improves efficiency and finds improved local solutions through efficient search of the parameter space.

When the GMM formulation is used to analyze multiple channels simultaneously, certain modifications need to be made for both statistical and computational considerations. Following [24], we fit a mixture model on the virtual masked data 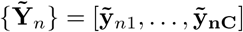, instead of the actual data {**Y***_n_*}. This allows the mixture model to use the localized nature of the data, which can dramatically reduce computations. Second, following [9, 24], we define the covariance structure in the GMM between channels to be 0. This step reduces the number of parameters to estimate in the covariance matrix to 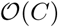 from 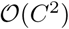.

This full model can be represented succinctly using a stick-breaking formulation of mixture weights [22] and a Normal-Wishart prior for each cluster and electrode. The assignment variable *z_n_* denotes which cluster the *n*th waveform is assigned to. Letting *k* define the cluster index, the full generative process of this model is

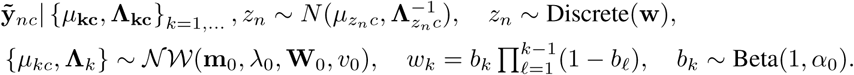

We note that if the mean and precision are known for a cluster, then inference for the optimal placement of a waveform exactly follows traditional GMM approaches. Second, we note that the stick-breaking formulation on **w** is such that the sum of the probabilities goes to 1 as *k* → ∞. While this has nice theoretical properties, this is a practically difficult representation, typically requiring the use of a Chinese Restaurant Process formulation [34] or adaptive methods such as Retrospective Sampling [10]. In practice, it is common to simply truncate the maximum value *K* at a high value.

We alternatively perform inference in this model via by adapting the Variational Bayesian (VB) split-merge approach of [21], which dynamically chooses this truncation level *K*, to utilize our coreset representation. The key idea of the split-merge approach is based on two moves. First, the merge, or cluster death, will combine two clusters if there is not sufficient statistical evidence to support distinct clusters. Vice versa, the split, or cluster birth, will take a cluster that represents an inhomogeneous waveform population and split it into multiple clusters (or neurons).

The first step in the mathematical formulation of this approach is to define approximate posterior forms. Letting Θ = {*μ_kc_*, Λ_*kc*_}_*k*=1,…,*K*,*c*=1,…,*C*_, this is given by

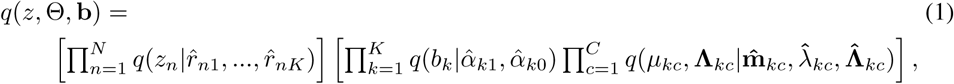

where 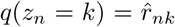, 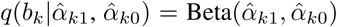, 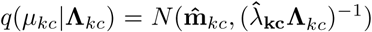, and 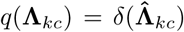, a delta function at a particular point. The use of a delta function on the precision is non-standard and breaks the mean-field formulation, but allows us to provide an option to enforce a minimum variance. Enforcing that the minimum cluster variance does not go below the known background variance can improve robustness is certain situations; however, it comes with an increase in computational costs, so the algorithm default is to not use a minimum variance. Precedence of this approximation in VB inference can be found in [21]. Because the DP has infinite mixture components, we implicitly assume that *q*(*z_n_* = *k*) = 0 ∀*k > K*. The variational parameters are learned to minimize the KL-divergence between the true posterior and the variational distribution, which maximizes the Evidence Lower Bound Objective (ELBO), given by

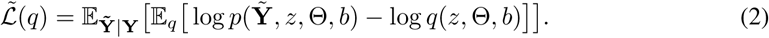

Compared to the typical ELBO used in [21], we have an expectation over 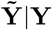 given by the masking approach. While this initially seems like it increases the complexity of the variational updates, the expectation over the mask will simply lead to a linear multiplicative factor when calculating updates. Hence, since the mask is typically 0, this allows great speedups by allowing sparse updates and reduces overfitting.

If the minimum variance constraint is set, then it is necessary to solve 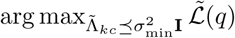 given the other parameters. Fortunately, a simple procedure can give the optimal solution. Succinctly, the MAP value from the standard variational update on the Wishart distribution is projected to the feasible set on its singular values. The projection is performed by taking the SVD, setting the projected singular values to the minimum of itself and 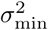, and reconstructing. It is straightforward to prove that this update is optimal on the feasible set. A detailed mathematical description of this update and how the standard updates from [21] change is given in the following section. This step gives a modest reduction in overfitting and improves stability at the expense of additional computation.

As discussed in previous sections, running on all data points can lead to slow and redundant computations, so we want to modify the existing VB structure to utilize computations only on the coreset. In mixture models of exponential families, the mean-field parameters for the approximate posterior are determined completely by additive sufficient statistics. Therefore, when working with a coreset, each representative point can store the sufficient statistics of its members that can easily be used when updating variational parameters. Importantly, despite using only a computationally-friendly small set of representative points, these representative points allow the sufficient statistics from each member point to be included in posterior estimates. Specifically, the *G* representative points, 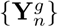, their masks, 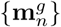, and their sufficient statistics, *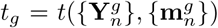* for the *G* representative points *n* = 1,…,*N_g_, g* = 1,…,*G* can be passed to the clustering algorithm to be used in the updates.

Once the ELBO is set up and modified to address masked data and representative points, there is a principled way to choose whether splitting or merging clusters is appropriate. We use the approach of [21], where following an update based off of the coreset, the algorithm proposes splits and merges to search for an optimal point. We give a high-level description of this in Algorithm S2, and further mathematical details in Section F.1.

### Algorithm S2 Overview of the DPMM procedure on a Coreset

Input: *G* sufficient statistics for each group, *t_g_*, *g* = 1, …, *G*

Initialize: number of clusters, *K*^(0)^, global parameters, *θ_k_*, and local parameters for each group and cluster, 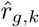, suff. stat. for each group and cluster, 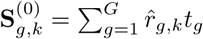

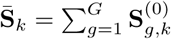

**for** i=1,… **do**

      Randomly pick a cluster *k*, split it into *K*′ + 1 clusters, yielding *K*^(*i*−1)^ + *K*′ clusters

      **for** g=1,…, G **do**

            Update 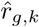 and 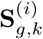

      **end for**

      Update *θ_k_*, for *k* = 1, · · ·, *K*^(*i*−1)^ + *K*′ using 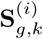

      Calculate 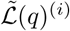

      Merge clusters if resulting 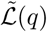 is lower than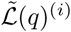. Let *K*^(*i*)^ ≤ *K*^(*i*−1)^ + *K*′ be the resulting number of clusters

**end for**

Choosing the hyperparameters in the DP-GMM will vary the performance of the algorithm. The default settings used in this algorithm are shown in Table S1, of which the most important are the hyperparameter for the stick-breaking parameter and the hyperparameters for the Normal-Wishart prior. The stick-breaking parameter was set to 1; however, similar performances were obtained from 10^−1^ to 10^1^. The hyperparameters for the Normal-Wishart were set such that the expected covariance of the cluster matches the background noise signal (e.g. **I**) with a non-informative mean.

### F.1 Additional Mathematical Details

Estimating the variational parameters is done such that they minimize the KL-divergence between the posterior distribution and variational distribution, which is equivalent to maximizing the evidence lower bound (ELBO). However, as the virtual data, 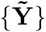, is not directly observed, the expected value of 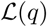 given {**Y***_n_*} is maximized. Accordingly, the following objective function is optimized:

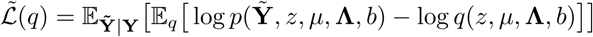

The local parameters given global parameters are updated as:

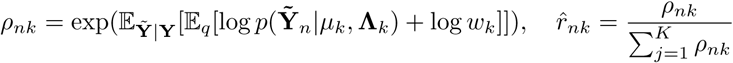

Given sufficient statistics, 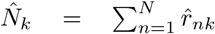, 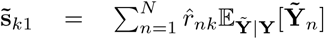, 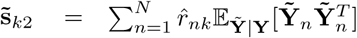, the global parameters are updated as following:

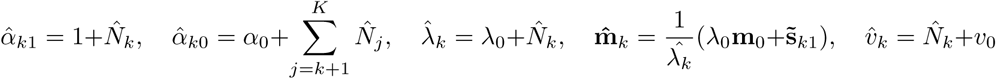

Updating 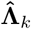 is a two-step process. 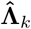 is first updated to maximize expected ELBO.

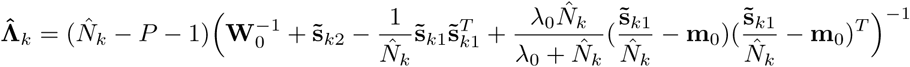

Then, let 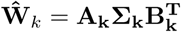,which is a singular valued ecomposition.The second update is

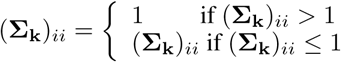

It ensures that the variance of each component is bigger than 1 as the variance is the sum of cluster variance and noise variance, which is one.

To further simplify the process, the independence of data across the channels can be assumed. Then, the covariance matrix of cluster has a block-diagonal shape and the cluster shape in each channel can be estimated separately using the above update. This is reasonable assumption as spatially whitened data is used.

The split and merge steps are conceptually the same as in [21], but adjusted to use the coreset.

## G Addditional details on the Divide and Conquer approach

Because of the method used to divide the data in the divide-and-conquer step, it is possible that the same neuron may have templates and clusters under different spatial subsets. Furthermore, the GMM approach may overcluster due to model mismatch (e.g. non-Gaussianity of the clusters). Therefore, it is necessary to merge the templates prior to the deconvolution step.

The templates are constructed as follows. After the clustering process, each spike **X***_n_* has been associated with one of *K* clusters, which is denoted by an assignment variable, *z_n_* ∊ {1,…,*K*}. Then, let 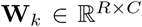 be defined as the mean of each cluster defined as *mean*({**X***_n_* |*z_n_* = *k*}), which is taken with respect to the original data rather than the virtual data distribution. In addition, a binary mask is defined for each cluster by 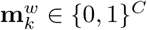, where

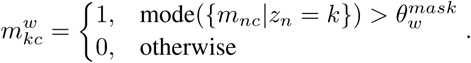

The mode on this continuous distribution is estimated by the peak of a kernel density estimate. 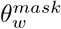 is set to 0.5. Given template masks, templates can be localized by considering channels with only non-zero mask entries. Template merging is performed based on two criteria, the angle and the amplitude of templates. When two templates are in fact the same neuron, the template waveforms should have a similar shape and their amplitudes should approximately match. To determine whether the waveforms have a similar shape, typically the cosine distance is used; however, the cosine distance is greatly affected when templates are not temporally aligned. Therefore, the angle” is calculated as the cosine distance on the best alignment between two templates. After the similarity check, an undirected graph can be created by considering each template as a node and constructed edges based off of similarity. Once the graph is obtained, strongly connected components are computed using Tarjan’s algorithm [42] to group templates. Then, new templates are created by taking the mean of each group. The pseudocode to perform this merging is shown in Algorithm S3.

## H Constructing a proxy to ground truth for a real dataset

It is notoriously difficult to obtain ground truth data for large-scale extracellular recordings of neural activity with MEAs. However, in rare cases, sufficiently strong anatomical and functional priors are available for the tissue under study and make it possible to hand-curate the outcome of a sorting pipeline, resulting in an acceptable partial proxy to ground truth on these data.

The data we used to construct such a proxy consisted of 30 minutes of extracellular recordings of the peripheral primate retina, obtained ex-vivo using a high-density 512-channel recording array [30]. During the half-hour recording, the retina was stimulated with spatio-temporal white noise in which a lattice of square pixels were updated randomly and independently of one another over time. The intensity of each display primary at each pixel location was chosen from a binary distribution at each refresh, yielding a stimulus with chromatic variation.

The reference set of spike times was manually assembled as follows. Events whose amplitude exceeded 4 times the RMS noise on each electrode were detected, aligned to the time of peak deviation from baseline using cubic spline interpolation for sub-sample alignment, and noise-whitened. Principal component analysis was performed on the collection of events detected on each electrode and the surrounding 6 electrodes, and the collection of events was clustered using a variation of the Ng-Jordan-Weiss spectral clustering algorithm [35, 52]. The hyper-parameters of the clustering algorithm were sweeped, resulting in a large collection of candidate neurons. Candidate neurons whose spike train exhibited significant violations of the refractory period were rejected.

Reference neurons were then identified on the basis of their light response properties. Anatomical and functional priors guarantee that classes of retinal ganglion cells in the primate retina tile the visual field uniformly, forming well-coordinated mosaics [11]. After calculating the spike-triggered average response of each neuron, cells were therefore separated in unique functional types, corresponding to the ON and OFF midget cells, ON and OFF parasol cells, ON and OFF upsilon cells, small bistratified cells, and polyaxonal amacrine cells. For each cell detected more than once in a mosaic, only the cell with the largest spike count was kept. All cells with physiological properties incompatible with the anatomical and functional priors of the primate retina were discarded from the analysis, resulting in a final collection of unique reference neurons of known cell types (n = 355), as well as a few neurons whose physiological properties were consistent with yet unreported cell types (n = 31).

### Algorithm S3 Template Merging

templates ← `mergeTemplates`({clusterAssignments*_i_*}*_i_*_=1,…_, {representativeWaveforms*_i_*}*_i_*_=1,…_)

Algorithmic Settings: Maximum shift allowance 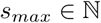, thresholds, *θ*_1_, *θ*_2_ ∊ (0, 1), number of clusters, *K*, temporal length of waveform, *R*, number of channels, *C*.

Input: clusterAssignments are given by {*z_n_*}*_n_*_=1,…,*N*_ ∊ {1, …, *K*} and representativeWaveforms are given by 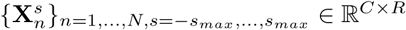, waveforms shifted by *s* from its center.

Output: templates

% Get templates with all shifts

**for** *k* = 1, …, *K* **do**

       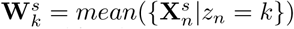

       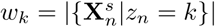

**end for**

Initialize: Undirected Graph *G* with *K* nodes and 0 Edges

**for** *k*_1_, *k*_2_ = 1, …, *K* **do**

       % shift that maximizes the cosine of two templates

       *s*′ = arg max*_s_ cosineDist* 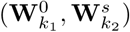

       **if** *cosineDist* 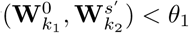 and 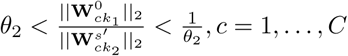 **then**

         Add edge on (*k*_1_, *k*_2_)

       **end if**

**end for**

Using strongly connected components on *G* [42], templates are grouped into *K*′ groups.

For *k* = 1, …, *K*′, new template 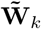, is the mean of properly shifted templates in each group weighted by *w_k_*.

Note that, by construction, this gold standard database includes a collection of trusted spike shapes and times, but we do not claim that this analysis captures every single true spikes from this recording; indeed, we expect that the dataset includes a number of unlabeled small or collided spikes.

## I Relationship to existing pipelines

Prior to the clustering stage, several important preprocessing steps take place. Filtering with wavelets [47] can improve sorting performance; however, we chose to use a bandpass Butterworth filter [29] because it is used in most academic and commercial implementations, such as the Plexon Offline Sorter^TM^.

The most common detection algorithm selects events that cross a threshold, with alignment to the threshold crossing or a peak [29]. Alternatively, [24] used multiple thresholds and temporal and spatial adjacency to improve detection, and aligned to the mean energy of a soft thresholded signal; [38] used wavelets to denoise and detect spikes. All these methods generally lead to many false positives that negatively affect the clustering step. Our pipeline improves on these issues by triaging false positives and addressing them in a later post-processing step (see Section C.1).

The clustering stage of the spike sorting problem has been discussed in several reviews [41, 29, 16, 39]. Spike sorting on dense MEAs has been explored via template matching [15], blind deconvolution [8, 37, 12], and clustering [24, 9]. Our approach is based on the Dirichlet process Gaussian mixture model (DP-GMM), which first introduced to the spike sorting problem by [48]. There have been significant improvements in efficient inference for the DP-GMM, and our work heavily utilizes the memoized approach of [21], but several alternatives exist, including online approaches [46], utilized in [8], and parallel computing [7].

Compared to prior work, we directly address collisions and outliers by excluding them from the clustering step with triaging, which enables significantly improved clustering performance. Furthermore, local minima are a critical problem in many clustering, template matching, and blind deconvolution approaches due to the non-convex nature of these approaches. While our model is also non-convex, our empirical results (not shown) demonstrated that adapting modern variational inference techniques improved both reliability and accuracy. These results match the conclusions of [21]. This is of crucial importance as the field moves towards millions of distinct waveforms, because common hand-sorting and hand-correction steps are intractable on such data.

There are a number of alternative approaches to the clustering step in the literature. Many of these approaches address the Gaussianity assumption on the shape of the clusters. One of the most common alternative approaches to a Gaussian mixture model formulation for clustering is a density based approach. This has been done in the form of superparamagnetic clustering [38], consensus clustering [14], unimodal clustering [31], spectral clustering, finding density peaks [40, 51], and the mean shift algorithm [20]. `MountainSort` [31] uses a non-parametric clustering step that assumes the clusters are unimodal in the sense that they have a single point of maximal density when projected onto any line. These approaches hold significant promise for dealing with non-Gaussian waveform clusters, and many of these clustering methods could be interchanged with our DPMM in our complete pipeline to alter the clustering step. However, as empirically demonstrated in our experiments, while these approaches can deliver improved performance on non-Gaussian clusters, on our metrics, for the datasets analyzed here, the average performance is below the proposed DPMM.

Recently, the `Kilosort` algorithm [36] was proposed to perform clustering and inference directly on the time series instead of using clustering. Collisions generally cause problems in the PCA and clustering space [12]; the matching pursuit approach of [36] sidesteps the clustering step entirely. In our pipeline, we address the collision problem with our triage-then-pursuit strategy; this lets us use fast clustering primitives to estimate templates, leading to significant scalability gains relative to full matching pursuit approaches. (Clustering approaches are also better able to handle uncertainty in template shape and spike assignments than greedy matching pursuit approaches.) After we have an appropriate clustering and waveform shape estimate, we use some of the methods proposed in [36] and [12] for the final collision-unmixing step. Other methods implicitly incorporate overlapping spike detection in their model training [8]; however, this approach has not yet been demonstrated to scale to large dense MEA datasets.

`Spyking Circus` [51] is another algorithm that aims to scale spike sorting to large MEA recordings by density estimation clustering of dimensionally reduced spikes by PCA, followed by template estimation and matching. Spikes are detected according to threshold crossings on each channel which can affect PCA projections negatively because of collisions and alignment issues. The algorithm is scaled to large recording sizes by sub-sampling spikes (data points), which can under represent units with lower firing rates. This method allows for parallelizing the computation of affinity matrices of data points which has lead to GPU and multi-threading implementations. However a great deal of post processing of results of each channel is needed that in practice renders the method time inefficient compared to `YASS` and `KiloSort`.

`JRClust` [23] follows a similar method as `Spyking Circus`. It does not use deconvolution to infer collisions. The method, however, addresses an issue of non-stationarity caused by noise and probe drift. Due to methodological similarity to `Spyking Circus`, it is expected that their performances are comparable. We hope to provide more detailed comparisons in the future.

## Footnotes

1 DARPA Neural Engineering System Design program BAA-16-09

2 Spyking Circus took over a day to process this dataset on a 6-core machine. The runtime of Mountainsort was not measured due to its relatively poor accuracy compared to Kilosort. Assuming linear scaling based on smaller-scale experiments, it is expected to take approximately 10 hours on a 6-core machine.

